# A comparison of long-read single-cell transcriptomic approaches

**DOI:** 10.1101/2025.07.03.662955

**Authors:** Anita L. Ahlert Scoones, Yuxuan Lan, Charlotte Utting, Lydia Pouncey, Ashleigh Lister, Sofia Kudasheva, Neelam Mehta, Naomi Irish, David Swarbreck, Karim Gharbi, Wilfried Haerty, Adam P Cribbs, David J. Wright, Iain C. Macaulay

## Abstract

Long-read sequencing enables the incorporation of isoform-level expression into single-cell transcriptomic studies, offering detail beyond those accessible with short-read methods. Although insightful, these approaches have typically been costly and yielded limited data for each individual cell. Recent advances in library preparation approaches and sequencing throughput have brought long-read single-cell studies closer to the mainstream.

Here, we present a comparative analysis of commercial approaches for single-cell long-read sequencing. We have performed parallel analyses of the same cDNA material, generated using the 10X genomics platform, on Illumina short-read, and PacBio and Oxford Nanopore long-read platforms. We also demonstrate the impact of CRISPR-based depletion of libraries, to remove highly expressed transcripts, prior to long-read sequencing in these experiments.

By analysing single-source cDNA libraries in parallel, we enable a direct comparison of each platform, evaluating standard metrics alongside concordance in clustering and cell type identification. While each approach generates usable gene and isoform expression data, we identify limitations common across platforms, primarily linked to cDNA synthesis inefficiencies and read filtering strategies. Our work demonstrates the increasing utility of single-cell long-read sequencing for isoform-resolved analyses, such as direct immunoglobulin chain reconstruction without additional amplification, and the detection of alternative splicing patterns across immune cell subtypes in CD45, a key gene for immune cell activation and differentiation. Our benchmarking of current platform options provides a foundation for researchers looking to adopt single-cell long-read sequencing into their transcriptomic studies, providing a framework for its integration into diverse biological questions.

## Introduction

Single-cell RNA-seq (scRNAseq) has transformed the study of multicellular systems by enabling transcriptomic profiling at cellular resolution. Recent advances in scRNAseq technologies have facilitated the parallel analysis of millions of cells, supporting the generation of comprehensive cell atlases at the organ and whole-organism scale ^1–4^.

As these technologies have advanced, there has been a near-universal focus on 3’ or 5’ transcript end counting to overcome the technical and economic constraints of high throughput single-cell measurements. Although this approach has been transformative, it inherently restricts the resolution of complex and biologically meaningful transcriptomic events - such as alternative splicing, gene fusions and single-nucleotide variants - across the full-length of the transcript.

Several studies have now demonstrated the value of long-read sequencing technologies, particularly Oxford Nanopore Technologies (ONT) and PacBio, in recovering full-length transcript sequences from both bulk ^5^ and single-cell datasets ^6–8^. These approaches enhance the detection of novel transcripts, landscapes of transcription, alternative splicing ^9^ and regulatory networks ^10^.

Increased scale of sequencing and improvement in read quality have made long-read sequencing of scRNAseq libraries a more realistic opportunity. In addition, concatemerization of cDNA molecules ^11^, and improvements with ordered concatemerization, termed MASseq ^12^ (now commercially supplied by PacBio as Kinnex), offer a novel approach to improve the output of the PacBio approach by as much 16-fold. Further modifications to library generation such as CRISPR-based depletion ^13^, allow more targeted sequencing, by eliminating highly expressed transcripts prior to sequencing, reducing consumption of sequencing capacity on uninformative reads.

Although these developments have expanded the practical utility of long-read scRNA-seq, many benchmarking studies predate these recent technical innovations. Here, we present a comparative analysis of long-read single-cell RNA-seq protocols on PacBio and ONT platforms, incorporating MAS-seq and targeted CRISPR-based depletion to remove highly expressed transcripts prior to sequencing. We assess these approaches in terms of read yield and informational content, as well as their utility for cell-type classification and isoform-level resolution within immune cell populations. Together, these analyses demonstrate the potential of long-read scRNA-seq to uncover isoform-level features in immune cells, providing a foundation for studying cell-state transitions and immune function with greater resolution.

## Results

### Biases in cDNA synthesis limit full-length transcript recovery in long-read scRNA-seq

We generated cDNA libraries using the 10X Genomics 3’ v3.1 platform and used the same cDNA pool as input into five different library preparation and sequencing approaches (Figure 1A). Our aim was to systematically compare how different sequencing platforms and library construction strategies, following manufacturer recommendations, enable isoform-level transcript resolution in single cells. Specifically, we evaluated long-read sequencing using the ONT PromethION (SQK-PCS111 kit) and the PacBio Revio and Sequel IIe platforms with the MAS-seq (now Kinnex) kit, alongside conventional short-read sequencing on the Illumina NovaSeq 6000.

**Figure 1.**
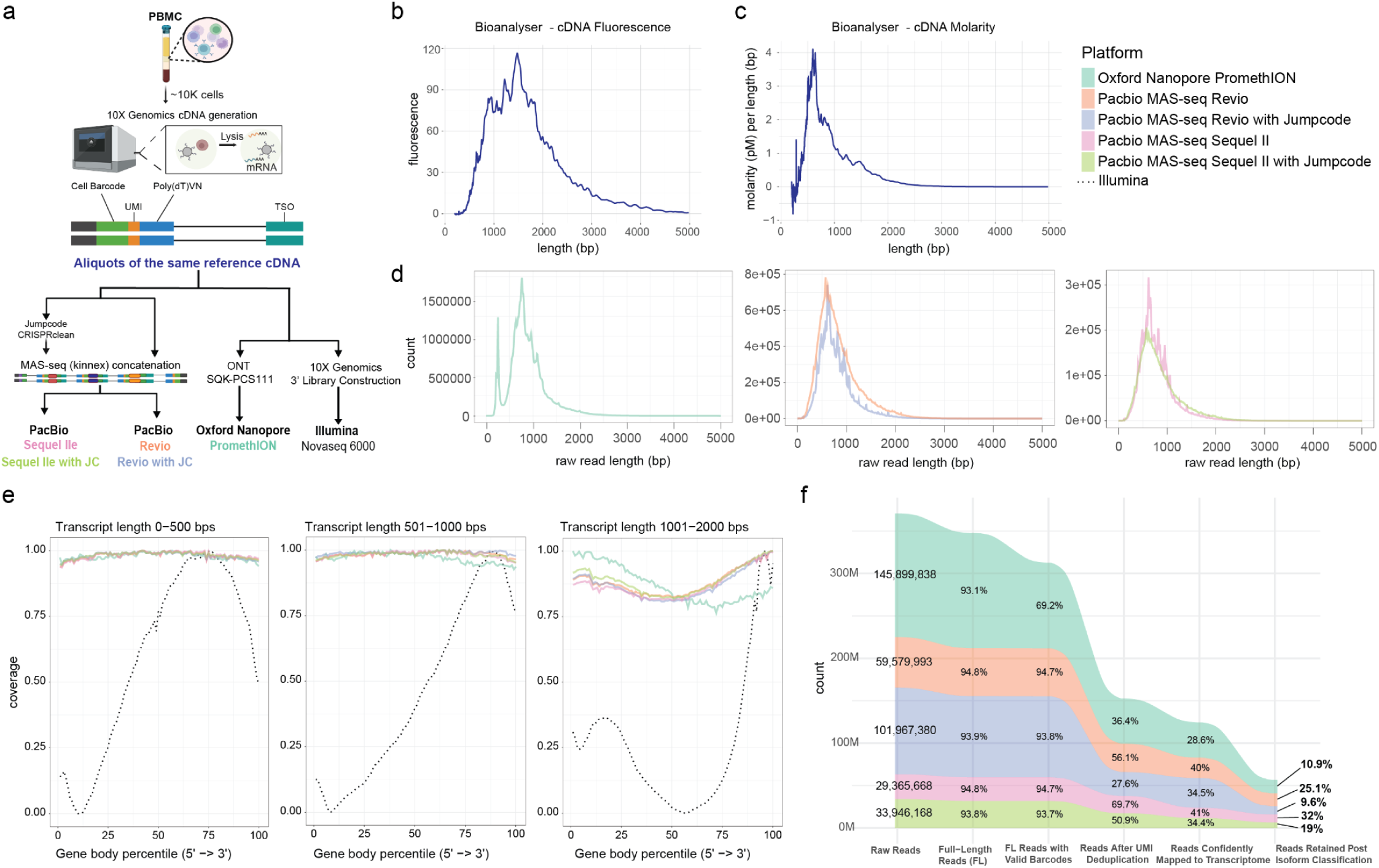
**Experimental workflow and long-read scRNA-seq data characteristics.** (a) Schematic of the experimental workflow. PBMC cDNA was generated using the 10X Genomics platform and split across multiple library preparation protocols, including PacBio MAS-seq (Revio and Sequel IIe), Oxford Nanopore PromethION, and Illumina (short-read) sequencing. Subsets of cDNA were treated with CRISPRclean depletion (Jumpcode) prior to MAS-seq preparation and PacBio sequencing. (b–c) Bioanalyser trace of the amplified cDNA showing (b) fluorescence intensity and (c) molarity distribution by fragment length. (d) Raw read length distributions for each long-read platform: Oxford Nanopore (left), PacBio Revio (centre), and PacBio Sequel IIe (right). (e) Gene body coverage of CDS from 5′ to 3′ for transcripts binned by length: 0–500 bp (left), 501–1000 bp (centre), and 1001–2000 bp (right), demonstrating read-through performance across platforms. Illumina (dashed line) shows 3′ bias due to library construction method. (f) Summary of read retention across pre-processing steps, including filtering for full-length reads, valid barcodes, UMI deduplication, confident mapping, and isoform classification. Values indicate absolute read counts and percentages retained at each stage.

The cDNA generated exhibited a typical length distribution for a 10X Genomics experiment. When visualised using fluorescence measurements on the y-axis (as is conventionally presented), the size distribution appeared to range from ∼500 bps to 2–3 kb (Figure 1B). However, it is important to note that fluorescence-based detection may overestimate the apparent length of the library, as shorter fragments incorporate less dye and therefore produce lower signal intensities. When molarity is instead plotted on the y-axis, it becomes clear that molecules longer than 1 kb are present at low abundance (Figure 1C). This pattern is reflected in the raw read length distributions across all sequencing platforms (Figure 1D, Supplementary Figure S3). ONT produced the longest reads, with a mean read length of 878 bp, while PacBio platforms showed slightly shorter read lengths, with Revio ranging from 785–839 bp and Sequel IIe from 782–842 bp.These findings highlight a key limitation of standard 10X Genomics cDNA, where the apparent length distribution overestimates molecule size, and most molecules are actually <800 bp, limiting full-length transcript recovery in long-read sequencing.

### Stratified gene body analysis reveals transcript length-dependent coverage across platforms

To evaluate gene body coverage across platforms, we categorised transcripts into defined length intervals rather than computing a global coverage distribution. This approach accounts for variability in transcript length and minimises biases that would otherwise distort coverage profiles. Differences in sequencing chemistry may lead to coverage dropout effects, particularly in longer transcripts, whereas shorter transcripts may exhibit higher terminal coverage due to fragmentation or incomplete cDNA synthesis.

By binning transcripts into three categories (0–500 bp, 500 bp–1 kb, >1 kb), we assessed how effectively long-read sequencing captures full-length transcripts across different gene sizes. Our analysis shows that reads surviving filtration steps enable full-length coverage of genes in the 500 bp–1 kb range across all long-read platforms, with far superior coverage of the transcript body and 5′ end compared to Illumina sequencing (Figure 1E).

We also compared the distribution of aligned bases across different genomic regions (intergenic, intronic, CDS, and UTR). As expected, most reads aligned to coding region of genes, with approximately 50% of aligned bases aligned to CDS across all platforms (Supplementary Figure S5). While the overall distribution of bases across genomic categories was similar between platforms, we observed that Jumpcode treatment reduced the proportion of bases aligning to intergenic and intronic regions, with a corresponding increase in UTR alignment. This is consistent with Jumpcode’s design to deplete highly abundant, uninformative transcripts and fragments, thereby enriching for reads mapping to more biologically relevant regions.

To further assess the extent to which reads-derived isoforms likely represent full-length transcripts, we compared the length distributions of detected isoforms with those of reference transcripts (Human_hg38_Gencode_v39 for PacBio, 10X genomics refdata-gex-GRCh38-2020-A for ONT) for both full splice match (FSM) and incomplete splice match (ISM) transcripts, as defined by SQANTI^14^. This analysis revealed that longer transcripts present in the reference were largely absent among the isoforms detected in our data, with transcripts longer than 1.5-2kb generally underrepresented across platforms (Supplementary Figure S4). Given that the same cDNA was used across all experiments, the observed differences in FSM transcript lengths between platforms suggest that the additional processing involved in MAS-seq library preparation may also contribute to biases toward shorter transcript recovery in long-read single-cell workflows (Supplementary Figures S6 and S7).

### Read retention is low across long-read scRNA-seq platforms but varies by preprocessing stage

While long-read sequencing offers advantages in transcript resolution and gene body coverage, data retention at each stage of processing is critical for maximising usable reads for downstream analyses. To assess this, we examined read loss at key preprocessing steps across platforms, evaluating how much sequencing data was ultimately retained and incorporated into the final expression matrices (Figure 1F). We processed PacBio and ONT data using their respective recommended scRNA-seq workflows, following standard analysis pipelines. This approach allows us to evaluate sequencing outputs as an end-user would obtain them and to assess how platform-specific processing influences read retention. While both workflows follow a similar overall structure, they differ in step order and incorporate platform-specific processing steps due to differences in sequencing chemistry.

Since the gene × cell and transcript × cell expression matrices form the foundation of downstream analyses, we examined read retention at each processing stage, tracking how much sequencing data was lost from the total raw reads to the final dataset used for expression analysis (Figure 1F). ONT generated the highest number of raw reads (145.9M), followed by Revio with Jumpcode (101.9M) and Sequel IIe (33.9M). Median read quality scores were substantially higher for PacBio (Q31–34) compared to ONT (Q13), reflecting known differences in sequencing accuracy across platforms.

Across all platforms, the percentage of full-length (FL) reads was consistently high (≥93%), indicating minimal data loss at this initial step. However, substantial read attrition occurred at later stages of processing. During the transition from FL reads to FL reads with valid barcodes, ONT retained only 69.2% of its total raw reads, compared to 93.7–94.8% in PacBio datasets (Figure 1F). This suggests that ONT’s higher sequencing error rate affects barcode correction, leading to early read attrition that is not observed in PacBio platforms.

The most substantial read loss across all platforms occurred by the UMI deduplication stage, where ONT retained only 36.4% of its original reads, indicating that nearly half of all ONT reads were removed during this step. PacBio platforms also experienced losses by this stage, with Revio retaining 56.1%, Sequel IIe retaining 60.7%, and Revio with Jumpcode treatment showing the largest reduction, retaining only 27.8% (Figure 1F). The steep decline in ONT suggests that error-prone UMIs contribute to increased filtering at this stage, whereas PacBio’s higher sequencing accuracy may lead to more stable read retention.

A notable difference between analyses is the incorporation of SQANTI3-based isoform classification and filtering in the PacBio workflow (*pigeon*), which removes low-confidence isoforms as a standard step, whereas the ONT workflow does not include an equivalent isoform filtering step by default. Previous studies (Lebrigand et al. 2020) and our own findings indicate that long-read single-cell datasets often contain a high proportion of unwanted artifacts, including template-switching by-products, ligation-derived adapter sequences, intrapriming events, and non-canonical splice site artifacts. These do not correspond to true cDNA structures yet consume a significant portion of sequencing throughput. As a result, ONT initially retained a far greater number of reads in the expression matrices, which could lead to an overestimation of isoform diversity. To ensure the analysis approaches were as comparable as possible between approaches, we applied default SQANTI3 filtering to the ONT dataset before generating the final count matrices for downstream analyses. After applying SQANTI3 filtering, the number of usable ONT reads dropped to 10.9% of the original dataset (15.85M reads), aligning closer with Revio (12%) and Sequel IIe (19%) (Figure 1F). This suggests that while ONT initially appeared to retain more usable reads, many of these were likely false isoforms that PacBio’s default pipeline would have already filtered out.

These analyses indicate that while 10X Genomics cDNA libraries introduce length bias, long-read sequencing achieves high gene body coverage across single-cell transcriptomes. Despite this, recovered transcript lengths consistently fall short of reference annotations, highlighting a need for protocol optimisation to enable true end-to-end transcript capture at single-cell resolution.

### Long-read platforms yield consistent barcode recovery but variable isoform detection at single cell resolution

To assess molecule recovery at single-cell resolution, we compared UMI counts per cell across sequencing platforms. Among long-read datasets, ONT and PacBio Revio recovered similar UMI counts per cell, differing by only ∼2%, while Sequel IIe detected substantially fewer UMIs (∼36% fewer) (Table 1). In comparison, Illumina short-read sequencing yielded the highest median UMI count per cell (5510), approximately 79% higher than ONT—the best-performing long-read platform for UMI recovery—indicating that short-read sequencing remains superior to long-read approaches in this metric (Figure 2A).

**Figure 2.**
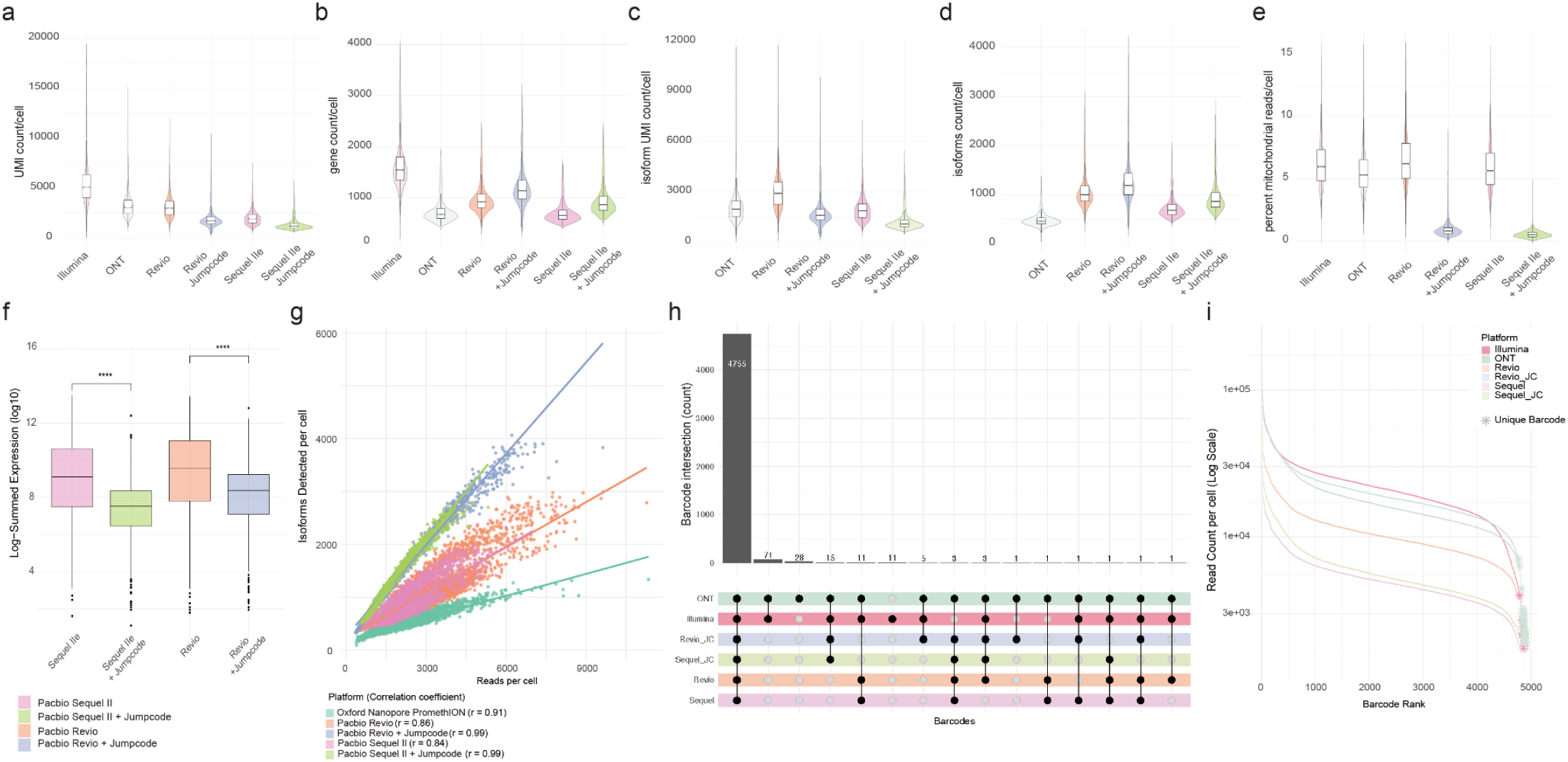
Comparing UMI recovery, isoform detection, and barcode overlap across scRNA-seq platforms. (a–e) Violin plots showing key QC metrics across platforms: (a) UMI counts per cell, (b) gene detected per cell, (c) isoform UMI counts per cell across long-read platforms, (d) isoforms detected per cell across long-read platforms, (e) percentage of mitochondrial reads per cell. (f) Summed expression (log₁-transformed) of targeted panel genes across Sequel IIe and Revio datasets, with and without Jumpcode treatment. Each box represents the distribution of log-summed expression values across targeted panel genes for untreated and Jumpcode-treated samples. Statistical significance was assessed using the Kolmogorov-Smirnov test and is indicated by asterisks (****, p < 0.0001). (g) Scatter plot showing the number of isoforms detected per cell as a function of read depth, lines represent linear regression fits. (h) UpSet plot showing the intersection of cell barcodes detected across sequencing platforms. Each row represents a platform, and each column of connected dots indicates a specific combination of platforms sharing the same barcodes. Bar heights reflect the number of barcodes per intersection. (i) Barcode rank plot comparing read count distributions across all platforms. Barcodes unique to a single platform are marked with bold asterisks.

**Table 1.** Summary of key metrics obtained from scRNA-seq data across different sequencing platforms. Metrics include the total number of cells analysed and retained post-quality control, RNA and gene detection per cell, mitochondrial read proportions, and isoform-level resolution.

| Seurat Object | Total Cells Analysed<br>(Original, % Retained) | Median RNA Counts per<br>Cell (Range) | Median Genes per<br>Cell (Range) | Mitochondrial Reads (% per<br>Cell, Range) | Median Isoform Counts per<br>Cell (Range) | Median Isoforms per<br>Cell (Range) |
| --- | --- | --- | --- | --- | --- | --- |
| Illumina | 4506 (4875, 92.4%) | 5102 (650-25071) | 1548 (369-3877) | 6 (0.8-15) | NA | NA |
| ONT | 4328 (4897, 88.4%) | 3076.5 (1203-14662) | 681 (450-1893) | 5.3 (1.4-15) | 1974 (382-11338) | 463 (88-1333) |
| Pacbio Revio MAS-seq | 4408 (4776, 92.3%) | 3006.5 (628-11571) | 925 (383-2376) | 6.2 (1.3-14.9) | 2912.5 (557-11270) | 1004 (368-2990) |
| Pacbio Revio MAS-seq with<br>Jumpcode | 4685 (4784, 97.9%) | 1720 (435-10203) | 1142 (378-3113) | 0.8 (0.1-8.8) | 1599 (360-9625) | 1191 (322-4063) |
| Pacbio Sequel Ile MAS-seq | 4212 (4773, 88.2%) | 1932 (736-7227) | 657 (457-1700) | 5.6 (1-14.7) | 1862.5 (647-7006) | 685 (425-1986) |
| Pacbio Sequel Ile MAS-seq<br>with Jumpcode | 4249 (4777, 88.9%) | 1183 (704-5636) | 869 (579-2362) | 0.5 (0-4.9) | 1084 (571-5321) | 867 (510-2828) |

While ONT yielded the highest UMI counts among long-read datasets, gene detection per cell was highest with PacBio MAS-seq combined with Jumpcode depletion on the Revio platform, recovering a median of 1,142 genes per cell—the highest among long-read libraries sequenced (Figure 2B, Table 1). Comparatively, ONT and Sequel IIe libraries detected approximately 40% and 43% fewer genes per cell on average. This trend was consistent at the isoform level, with the Revio MAS-seq Jumpcode library demonstrating the strongest performance, both in the median number of isoform UMIs (2,912) and isoforms detected per cell (1,004) (Figures 2C-D, Table 1).

For Jumpcode-treated libraries, we anticipated a reduction in mitochondrial transcript content, as the kit is designed to deplete highly expressed, often uninformative genes, thereby preserving sequencing capacity molecules of greater biological interest. Our analysis confirmed this, with mitochondrial transcripts decreasing by ∼87-91% following Jumpcode treatment, from from ∼5–6% in untreated samples to 0.6-0.8% in treated libraries (Figure 2E). Consistent with this, treatment with Jumpcode improved per-cell gene detection on both MAS-seq libraries, increasing by 24% on Revio and 32% on Sequel IIe, suggesting that depletion of highly abundant transcripts enhanced transcriptome complexity.

To further evaluate the effectiveness of Jumpcode treatment in depleting target transcripts, we compared the summed expression of all genes from the Jumpcode panel across untreated and Jumpcode-treated samples sequenced on both the Sequel IIe and Revio platforms. Jumpcode treatment resulted in a substantial and statistically significant reduction in panel gene expression across both platforms (Figure 2F). In Sequel IIe samples, Jumpcode reduced panel gene expression by approximately 2.9-fold compared to untreated controls (log₂ fold change = 1.55; Kolmogorov-Smirnov D = 0.427, p = 6.18 × 10⁻²⁷). Similarly, in Revio samples, a ∼2.3-fold reduction was observed following treatment (log₂ fold change = 1.21; D = 0.340, p = 2.84 × 10⁻¹⁷). To assess whether Jumpcode depletion impacted coverage of untargetted transcripts (i.e., transcripts not included in the panel), we examined distributional shifts and found minimal differences following treatment (D = 0.092 for Sequel IIe and D = 0.063 for Revio), suggesting depletion was highly specific to the targeted gene set without broadly affecting non-target transcripts. ECDF plots illustrating these distributional differences for panel and non-panel genes are provided in Supplementary Figure S1.

We next examined the relationship between sequencing depth and isoform detection across platforms (Figure 2G). Across all datasets, we observed a strong positive correlation between the number of reads and isoforms detected per cell. This relationship was most pronounced in Jumpcode-treated libraries, with both PacBio Revio and Sequel IIe exhibiting near-perfect correlations (r = 0.99), indicating highly efficient and consistent isoform recovery following depletion, reinforcing the utility of targeted depletion for isoform-level resolution in long-read protocols.

Given that all libraries contain the same cells sequenced across platforms, we next evaluated whether cell barcodes were detected consistently across technologies despite platform-dependent differences in molecular recovery. We observed high concordance in barcode detection across platforms, with most cell barcodes consistently recovered irrespective of technology (Figure 2H). While a small number of barcodes were detected exclusively in Illumina (n = 11) and ONT (n = 28) datasets (Figure 2H); these cells exhibited markedly lower UMI counts and were among the poorest-performing barcodes (Figures 2I). Their limited expression profiles suggest these barcodes represent low-quality cells rather than genuinely platform-specific captures.

In summary, combining Revio MAS-seq with Jumpcode depletion yielded the most comprehensive isoform recovery across platforms tested. Although ONT achieved marginally higher UMI recovery among the long-read datasets, this did not translate to greater gene or isoform detection per cell. Moreover, these results show that applying targeted transcript depletion enhanced transcriptome complexity at isoform resolution, supporting their utility in single-cell long-read sequencing workflows.

### Long-read scRNA-seq refines cluster annotation and enhances subpopulation resolution

To enable meaningful comparisons between platforms, all datasets underwent consistent quality control and were processed independently using the same parameters. Cells were visualised using Uniform Manifold Approximation and Projection (UMAP) at a fixed resolution and annotated based on canonical marker expression, supported by automated cell type prediction tools. Each dataset was annotated in isolation, without reference to labels from other platforms. We identified the same broad immune cell types across all six datasets, consistent with the composition of PBMCs, including T cells (CD4+, CD8+), B cells, natural killer (NK) cells, monocytes (CD14+, CD16+), dendritic cells (DCs) and platelets.

To evaluate annotation consistency, we compared cell type assignments to those derived from Illumina short-read data, and assessed the added impact of incorporating isoform-level information on cell type resolution. Notably, B cells and NK cells were the most reliably annotated across all platforms, with minimal variation in assigned labels (Supplementary Figure S2). These populations exhibited high concordance with short-read -based annotations and were recovered at similar proportions, reflecting their distinct transcriptional profiles and robustness to differences in platform sensitivity. In contrast, the annotation of T cell subsets showed greater variability, particularly CD8+ sub-populations. This may reflect the finer transcriptional distinctions among T cell states, and highlights differences in how each platform resolves transcript complexity and gene identity.

To assess how isoform-level information influenced cell-type resolution, we compared clustering and annotation across datasets using gene-level, isoform-level and integrated data (Figure 3). Major immune cell types remained consistent across analysis modes, but isoform-level data provided improved resolution of specific populations and greater separation between closely related T cell subsets. Sankey plots revealed that while most cells retained consistent annotations across gene- and isoform-level assignments, a subset of cells shifted labels—most frequently between closely related T cell states (Figure 3D, H), suggesting isoform data captured transcript features not evident at the gene level, refining cell identity in ambiguous regions. Isoform-aware annotation resulted in reclassification across all platforms (Figure 3). These findings indicate that isoform-level analysis adds resolution beyond gene expression alone, particularly for fine-grained distinctions among populations, for instance, T cells.

**Figure 3.**
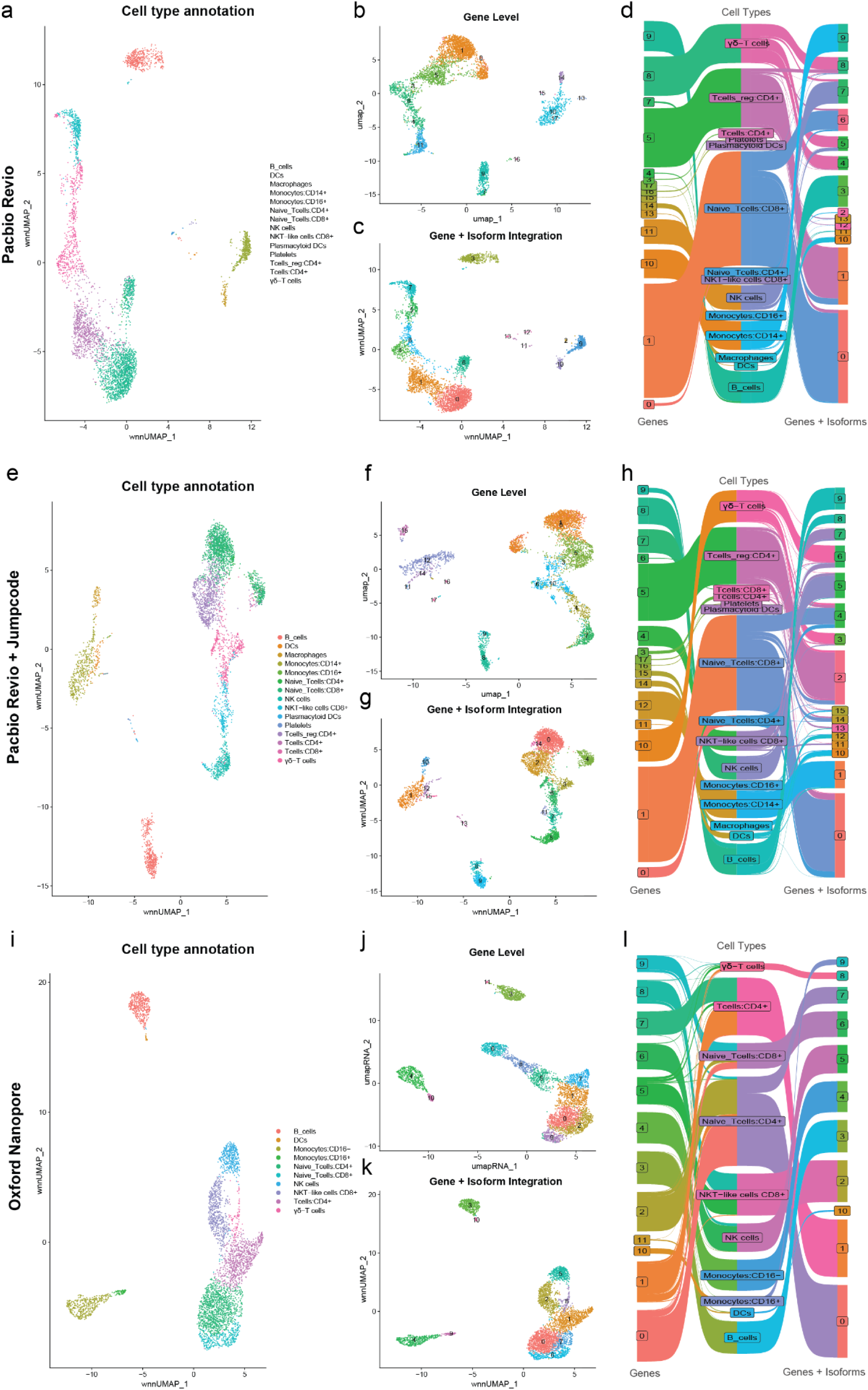
Gene- and isoform-level clustering and cell type annotation across long-read platforms. UMAP projections of PacBio MAS-seq on Revio (a–c), PacBio MAS-seq on Revio with Jumpcode (e–g), and ONT (i–k) datasets, based on cell type annotation labels (a e, i), gene-level clusters (b, f, j), and integrated gene and isoform expression clusters (c, g, k). Sankey plots (d, h, l) show corresponding cell type annotations derived from gene-level (left) compared to gene with isoform-level (right) quantification, where central labels indicating final cell type assignments.

### Direct detection of immune repertoires without targeted amplification and improved resolution of γδ T cells with isoform-aware scRNA-seq

Single cell clustering of long read data captured the same cell types at similar proportions, with the notable exception of a population of γδ T-cells that were not detected in conventional 10X-Illumina sequencing (Figure 4A). To explore this further, we subsequently used Trust4 ^15^ to perform immune repertoire analysis of the long-read data from all samples. Trust4 reconstructs recombined TCR and BCR sequences directly from RNA-seq data, using full-length transcript alignments to infer V(D)J gene usage and CDR3 sequences.

**Figure 4.**
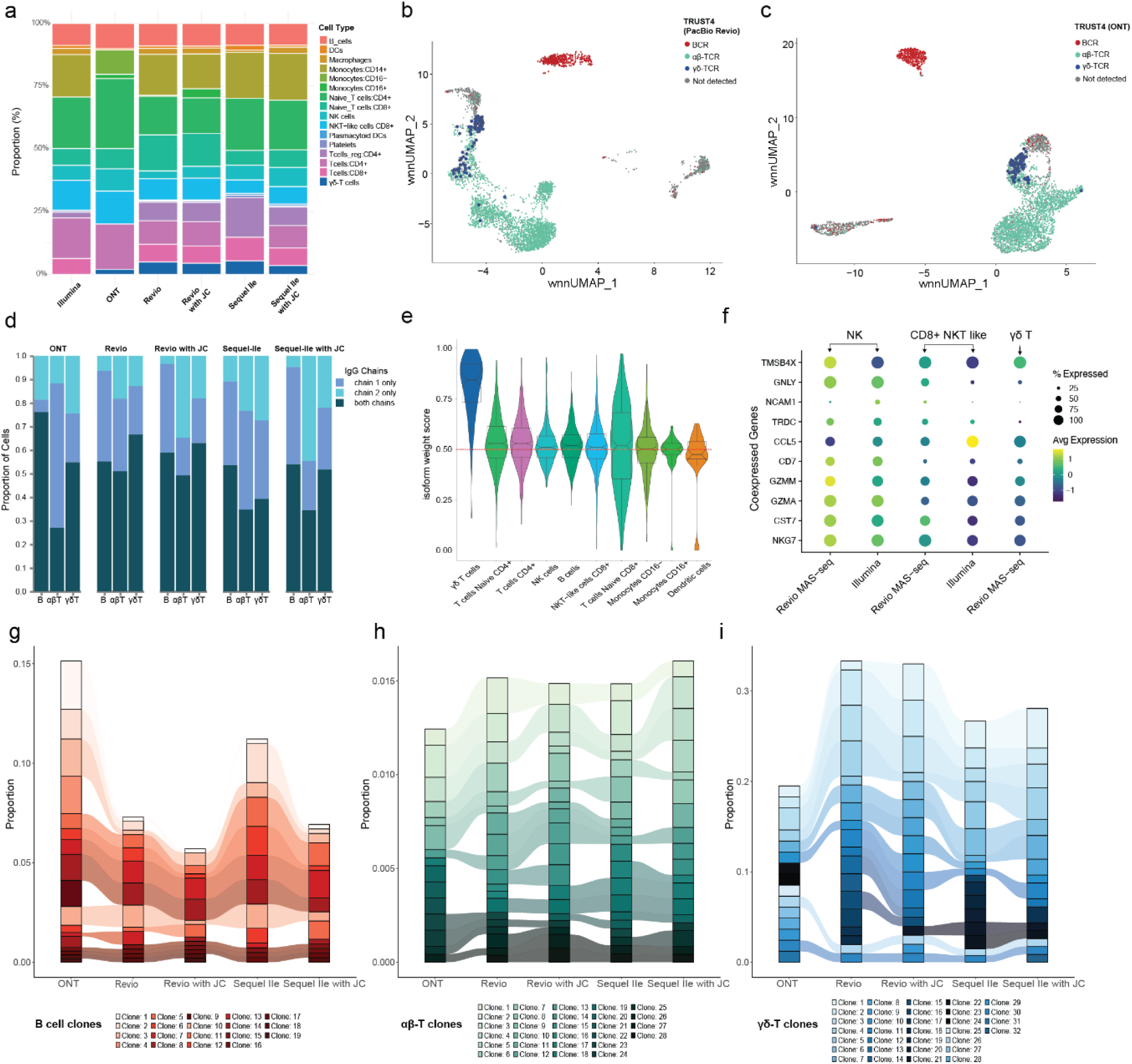
Long-read sequencing enables isoform-level resolution to improve annotation and characterisation of immune populations. (a) Cell type annotation proportions across platforms and chemistries has consistent recovery of major immune populations, with the exception of γδ T cells (b, c) WNN UMAPs coloured by TRUST4 immune receptor calls for Revio (b) and ONT (c). (d) Distribution of Ig and TCR chain recovery across immune subsets, demonstrating recovery of paired immune receptor chains with long-read sequencing. (e) Isoform weight scores across cell types, highlighting high isoform contribution in γδ T cell clustering. The red dashed line indicates equal contribution of isoforms and genes to cluster identity. (f) Feature expression dot plot showing co-expression of γδ T cell and NK cell markers across Illumina and Revio MAS-seq libraries. (g-i) B cell (g), αβ T cell (h), and γδ T cell (i) clonal distributions across long-read platforms, generated using scRepertoire, reveal platform-specific differences in resolving clonal architecture across immune subsets.

In conventional single-cell analyses, immune receptor profiling typically requires preparation of dedicated sublibraries for either class of receptor through targeted 5’ amplification, generating separate libraries for gene expression and αβ-TCR/BCR detection. Commercial approaches are available for mouse and human ɑβ-TCR and BCR sequencing, but a custom approach must be employed for γδ-TCR sequencing^16^, or immune profiling in non-model organisms. We demonstrate that long-read sequencing of single-cell libraries can uniquely enable detection of all classes of recombined TCR/BCR molecules without additional library preparation, including both ɑβ and γδ T-cells/TCRs (Figure 4 B-C).

We assessed the detection of full-length paired chains in B, αβ T, and γδ T cells (Figure 4D) to evaluate immune receptor recovery across platforms. Across all datasets, a substantial proportion of cells had both chains recovered, with the highest proportions observed in ONT and MAS-seq Revio libraries. Notably, full γδ TCR pairing was reliably achieved in both Revio libraries, highlighting the ability of high-fidelity long-read platforms to capture complete receptor sequences without enrichment. Sequel IIe libraries showed lower overall pairing rates, though performance improved with Jumpcode depletion. These results reinforce the potential of long-read RNA-seq for immune repertoire profiling.

To quantify the relative contribution of gene- and isoform-level data to cluster identity, we examined WNN modality weights, which reflect the information content of each modality at the single-cell level. Values near 1 indicate that isoform-level features are more informative for a given cell, whereas values near 0 reflect dominance of gene-level information. We visualised the distribution of isoform modality weights across WNN clusters annotated by cell type (Figure 4E). γδ T cells showed the highest isoform modality weights, indicating that isoform-level data contributed disproportionately to their identification. In contrast, monocytes and dendritic cells showed more balanced or gene-dominated weights, suggesting that isoform-level resolution provided less additional information for these cell types. These results highlight cell-type-specific differences in the added value of isoform data, particularly in lineages where transcript structure may be more functionally relevant or discriminative.

Prompted by the high isoform weight scores in γδ T cell clusters in our long-read data — not distinguishable as γδ T cells in our matched Illumina short-read dataset — we explored why gene-level signals alone may have been insufficient to resolve them. γδ T cells are a rare lymphocyte population (1–5% of circulating T cells^17^) that adopt an alternative TCR architecture, expressing TCR γ and δ chains (e.g., TRDC) rather than the αβ chains predominant in conventional CD3+ T cells. This grants them the ability to detect a wider array of antigens, including endogenous and exogenous non-peptidic molecules and stress-induced antigens, bypassing the need for MHC mediation^18^. Bridging innate and adaptive immunity, γδ T cells exhibit cytotoxic capabilities and cytokine profiles that closely parallel those of NK cells, reflecting their shared roles in early immune responses. In line with this, we found substantial overlap between gene-level markers for γδ T and NK cells in our dataset. For example, TRDC, a canonical γδ T cell marker, was robustly expressed in NK cells, which co-expressed CD56 (NCAM1), NKG7, GZMA, GNLY, CST7, and CCL5 (Figure 4F). Notably, TRDC expression in NK cells has been reported previously ^19^, although its functional relevance remains unclear. Consistent with this overlap, when examining matched cell barcodes in our Illumina short-read dataset, these same cells were annotated predominantly as NK or NKT-like cells (Supplementary Figure S8), highlighting the limitations of gene-level classification in resolving γδ T cells within standard scRNA-seq data. Our data suggest that γδ T cells—identified exclusively in long-read data through TRUST4 TCR reconstruction—can be more accurately distinguished from NK cells in scRNA-seq through long-read sequencing. These findings illustrate how isoform-level resolution can disentangle overlapping transcriptional signatures and improve annotation of closely related but functionally distinct immune subsets. This has important implications for immunology, particularly in efforts to characterise rare or unconventional immune populations, trace their lineage relationships, and understand their distinct effector functions. Specifically, in contexts such as tumour immunology or adoptive cell therapy development, where γδ T and NK cells are of interest for their cytotoxic potential ^20,21^, improved isoform-level resolution may facilitate mechanistic insights and support therapeutic targeting of these populations.

### Platform-dependent differences in clonal architecture captured by long-read scRNA-seq

To further assess the ability of long-read sequencing platforms to reconstruct immune repertoires, we used scRepertoire to assign clonotypes based on strict sequence identity and compared clonal distributions across datasets. Since immune receptor analysis is a key application of full-length transcript sequencing, this allowed us to evaluate how well each platform captures clonal architecture across major lymphocyte subsets. TCR and BCR sequences were analysed separately and grouped into B cells, αβ T cells, and γδ T cells (Figure 4G–I), where clonotype similarity was quantified using Morisita’s index to evaluate overlap across datasets.

B cell clonotypes showed the most consistent patterns, with dominant clones broadly shared across all datasets, as visualised in the riverplot (Figure 4G). While ONT showed a higher proportion of its top B cell clone relative to other technologies, which may reflect its higher recovery of paired BCR chains (Figure 4D), the overall BCR repertoire appeared relatively stable across methods, with similar rank and diversity profiles.

For αβ T cells, the PacBio MAS-seq Revio library had the highest number of unique clones identified, consistent with its greater recovery of fully paired αβ TCR chains. While distributions were broadly similar across platforms, ONT detected fewer total clones, and its repertoire appeared more skewed toward top-ranked clonotypes. In contrast, PacBio Revio and Revio with Jumpcode libraries exhibited smoother clone distributions, with greater retention of intermediate-abundance clonotypes (Figure 4H), suggesting improved resolution of subdominant T cell expansions.

γδ T cell clonotype distributions were more variable across platforms and skewed towards a smaller number of dominant clones (Figure 4I). This likely reflects a combination of lower γδ T cell abundance in the input sample and variability in paired-chain recovery, which limits the number of assignable clonotypes. Interestingly, both Jumpcode libraries retained a greater proportion of top γδ clones compared to their untreated counterpart, suggesting that depletion may help preserve dominant clonotypes by reducing background reads.

Overall, this analysis demonstrates that long-read single-cell RNA-seq enables reconstruction of clonal immune architecture across multiple lymphocyte lineages without dedicated receptor enrichment. The differences observed in clonal distributions here reflect technical variation in full-length receptor recovery rather than biological differences, as all datasets derive from the same input cell population.

### Cell-type-specific CD45 splicing captured by isoform-resolved scRNA-seq

To explore the utility of single-cell long-read sequencing in immunology, we leveraged TRUST4 classifications to separate reads by immune cell type (B cells, αβ T cells, and γδ T cells) for read-level splicing analysis ^15^. This allowed for a direct, read-level analysis of splicing patterns across immune cell types and study of base modifications. We focused on PTPRC (CD45), a regulator of white blood cell activation and differentiation, where exons 4, 5, and 6 are subject to alternative splicing to produce isoforms such as CD45RA, CD45RO, and CD45RB (Figure 5A). These isoforms influence TCR signalling thresholds and the activation states of naive, memory, and effector T cells, with evidence supporting conserved, cell-type-specific splicing patterns across immune lineages ^22^. The evidence for distinct functional roles across CD45 isoforms remains inconclusive, but there is strong support for conserved cell-type and differentiation-stage specific expression patterns in immune cell development ^23–25^. Moreover, dysregulation of CD45 splicing has been implicated in autoimmunity and haematological malignancies, yet the precise landscape of isoform expression at single-cell resolution has remained largely unexplored. ^26–28^. Our analysis revealed that B cells predominantly expressed isoforms retaining exons 4–6, corresponding to CD45RABC, whereas T cells primarily expressed shorter isoforms lacking these exons (Figure 5B). In contrast, gene-level expression of PTPRC appeared uniform across immune cell types (Figure 5C), highlighting the limitations of gene abundance alone in resolving biologically relevant splicing differences revealed by isoform-level analysis. Isoform-resolved profiling at single-cell resolution thus provides a more informative view of post-transcriptional regulation in immune cells and enables the study of splicing patterns in immunology.

**Figure 5.**
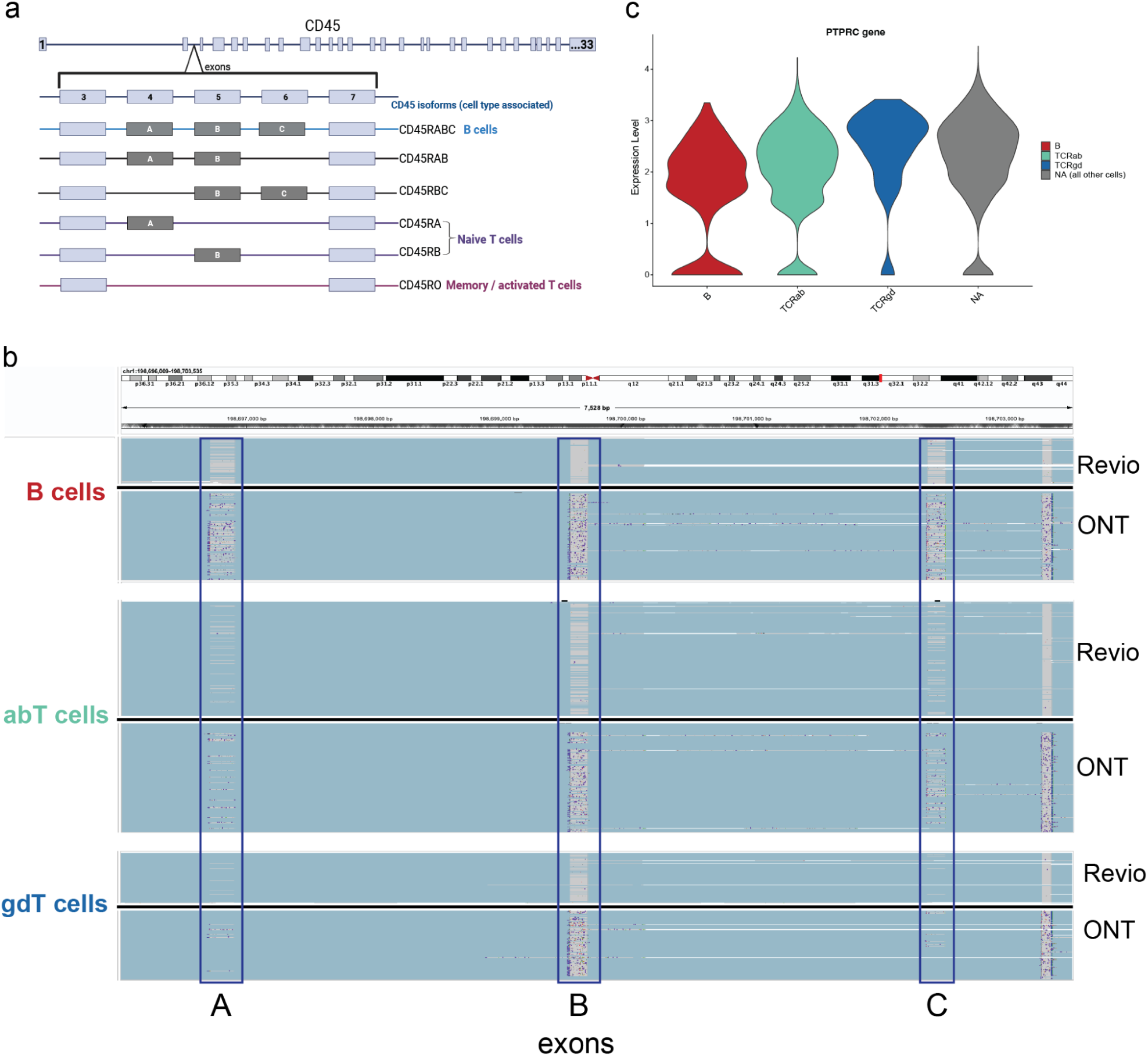
Single-cell long-read sequencing resolves PTPRC (CD45) isoform diversity across immune cell types. (a) Principal splice variants of CD45, illustrating alternative exon inclusion patterns that generate functionally distinct isoforms. (b) Read alignment tracks for the PTPRC region spanning exons 4–7 in B cells, αβ T cells, and γδ T cells using PacBio Revio (top) and ONT (bottom). Individual read alignments are shown, with indels indicated in purple. (c) Violin gene-level expression of PTPRC across B cells, αβ T cells, and γδ T cells appears broadly uniform, masking the diversity in alternatively spliced isoforms revealed by isoform-level analysis (Revio data shown).

## Discussion

This study provides a comparative analysis of single-cell long-read transcriptomics workflows, evaluating platform-specific wet-lab protocols and recommended bioinformatics pipelines under realistic usage conditions. Our analyses demonstrate that both platforms can recover complex transcript isoforms across diverse immune cell types, with comparable performance in barcode recovery, cell type identification, and overall per-cell resolution. However, we also highlight key differences between platforms that influenced downstream interpretation, shaping the granularity of isoform detection and the resolution of cellular subtypes.

To objectively evaluate platform performance, we analysed each dataset independently, treating each experiment in isolation while ensuring consistent parameters throughout sample processing and data interpretation. Across all datasets, the addition of isoform-level information restructured cell type clustering, often improving the resolution of closely related subsets, as exemplified by the improved identification of γδ T cells in long-read data relative to short-read sequencing. We also show that immune receptor repertoires can be reconstructed from standard gene expression libraries, extending the utility of long-read sequencing beyond dedicated immune profiling workflows.

While long-read approaches capture significantly more full-length transcripts than traditional short-read methods, transcript completeness declines sharply above ∼1 kb, reflecting a persistent limitation compounded by single-cell-specific artefacts such as mispriming and overamplification. Together, these factors highlight the need for optimised cDNA synthesis strategies prior to library preparation to reduce truncation and improve transcript recovery. In this study, while we did not deviate from manufacturer-recommended protocols to enable fair comparisons across platforms, modifications to cDNA synthesis workflows—including reverse transcription, amplification, and purification steps—represent a feasible and actively tested avenue for improving the recovery of longer transcripts. We demonstrated that CRISPR-based depletion of highly abundant transcripts, such as mitochondrial and ribosomal RNA, substantially improved the recovery of informative features. By redistributing sequencing capacity, depletion increased both gene and isoform detection. In the current landscape of limited throughput, targeted strategies such as transcript depletion or enrichment is proving the most effective means of enhancing coverage of transcriptional heterogeneity ^29^, particularly when specific biological processes or gene families are of interest.

These experimental challenges were mirrored by limitations in data interpretation. Unlike gene-level analyses, isoform quantification requires precise inference of transcript boundaries, splice junctions, and exon connectivity, making analyses highly sensitive to technical artefacts ^30^. Current pipelines, largely adapted from bulk data, do not adequately account for key sources of noise—including internal priming, non-canonical junctions, and poly(A) tail miscalls that are prevalent in scRNA-seq ^31,32^—which not only convolute interpretation of results but also leads to substantial data loss during pre-processing. Moreover, isoform detection and quantification, particularly for rare or alternatively spliced transcripts, require substantially higher sequencing depths than gene-level quantification due to the increased complexity of transcript structures. Our analysis revealed the substantial extent to which reads are filtered during early processing steps when following the recommended pre-processing workflows for each platform. Currently, artefacts such as chimeric, truncated, or misprimed molecules, and the inherent sparsity and variability of single-cell datasets remain unaddressed, limiting confidence in accurate isoform assignment. As the field matures, the development of platform-aware, artefact-sensitive pipelines will be essential to enable robust isoform-level analyses in single-cell contexts.

The selection of the most appropriate single-cell long-read sequencing approach to incorporate into transcriptomic studies, will also require considerations beyond data yield and quality. This includes costs, laboratory and bioinformatic staff time, training and the laboratory facilities available. In line with this, we have approximated the current processing time from cell isolation to completion of sequencing for each of the platforms implemented in this study (Supplementary Figure S9), though we acknowledge this will vary significantly between facilities and with continued development in the field. Since the completion of this work, further developments in long-read single-cell transcriptomics workflows have continued to advance. At the time of writing, the MAS-seq protocol, rebranded as Kinnex, now supports sample multiplexing and 5′-end compatibility, broadening its applicability across single-cell preparations. Similarly, the Oxford Nanopore libraries generated here using the SQK-PCS111 kit on R9 flow cells have been superseded by updated chemistries with improved read accuracy and yield. These developments reflect the rapid pace of progress in the field and a growing trend towards platform interoperability. Emerging combinations, such as pairing long-read sequencing with other single-cell approaches, point towards the potential to scale full-length isoform capture to tens of thousands of cells in the near future.

Despite these advances, it is important to recognise that the prevailing emphasis on cell throughput continues to constrain per-cell resolution. This trade-off is particularly limiting in biological contexts where low-abundance transcripts and subtle isoform changes can have functional significance, such as in lineage commitment and cell-state transitions ^33^. Consistent with the findings of others, our results indicate that long-read sequencing is well positioned to address these challenges. Realising this potential, however, will require a deliberate shift in strategy prioritising deeper sequencing of fewer cells and developing informatic tools capable of extracting robust isoform-level insights from single-cell data.

Overall, this study demonstrates that single-cell long-read sequencing provides practical and actionable isoform-level resolution for refining cell type annotation and repertoire analysis across immune cell types. Researchers seeking to investigate transcript diversity, splicing patterns, and clonal architecture should consider long-read approaches while balancing current throughput and cost constraints. As sequencing chemistries and informatic tools continue to advance, single-cell long-read sequencing will become increasingly integral to characterising transcriptional heterogeneity and post-transcriptional regulation, offering deeper insights into immune cell function in health and disease.

## Methods

### Single-cell cDNA preparation

Human Peripheral Blood Mononuclear Cells (PBMCs) from StemCell Technologies (Cat. No: 70025.1) were processed according to the manufacturer’s handling protocol. Cell viability and concentration were measured using a Countess II FL Automated Cell Counter (Invitrogen), and the suspension was diluted to approximately 1.18 × 10⁶ cells/mL. Single-cell partitioning and reverse transcription were performed using the Chromium Next GEM Single Cell 3’ GEM, Library & Gel Bead Kit v3.1 (PN-1000269) from 10X Genomics, following the manufacturer’s protocol (Revision E). Cells were loaded targeting a recovery of 9,000 cells per sample, and cDNA was amplified with 11 PCR cycles. The size distribution and concentration of amplified cDNA were assessed using a Bioanalyzer (Agilent Technologies) and a Qubit 1.0 fluorometer with the High Sensitivity DNA Qubit assay (Invitrogen). Although cDNA was generated from two PBMC samples, sufficient material for library preparation and sequencing across all platforms was only available from Sample 1; therefore, all analyses in this study were conducted using data from this sample.

### 10X 3’ Library preparation and sequencing

Aliquots of amplified cDNA from PBMC Samples 1 and 2 (10 μL) were used to prepare conventional Illumina libraries using the 10X Genomics Chromium Next GEM Single Cell 3’ v3.1 protocol (Revision E). 3’ Gene Expression Dual Index Library Construction was performed with the Dual Index Kit TT Set A (PN-1000215, 10X Genomics), using 13 PCR cycles for library amplification. Libraries were equimolarly pooled and sequenced on an Illumina NovaSeq 6000 instrument using an SP v1.5 flow cell, generating 100 bp paired-end reads across two lanes.

### Data processing and bioinformatic analysis of Illumina short-read data

Illumina single cell RNA-seq data were processed using Cell Ranger v6.0.1 against the GRCh38-2020-A reference transcriptome. The resulting feature-barcode matrices, containing gene expression counts, were further analysed in RStudio (version 4.3.2) using the Seurat R package (v5.1.0) ^34^.

Gene expression matrices were used to generate Seurat objects. Quality control thresholds were applied to exclude cells with unusually low or high numbers of RNA features (10th and 99th percentiles, respectively) and elevated mitochondrial read proportions (>15%).

To account for batch effects, PBMC samples were integrated using canonical correlation analysis (CCA) via Seurat’s IntegrateLayers function. Following integration, global-scaling normalisation was performed, and features were scaled. Dimensionality reduction was conducted using PCA and UMAP based on the first thirty principal components, with clustering performed at a resolution of 0.8 to identify cellular populations.

Cell type annotations were assigned based on canonical marker gene expression cross-validated using the automated annotation tools ScType ^35^ and SingleR ^36^, followed by manual refinement.

### Oxford Nanopore library preparation and sequencing

An aliquot of amplified cDNA from PBMC Sample 1 (10 ng) was used as input for library preparation, following the Oxford Nanopore Single-cell Sequencing on PromethION protocol (Revision C) with the SQK-PCS111 kit, using 14 PCR cycles for library amplification. Libraries were loaded at 25 fmol and sequenced on a PromethION P24 instrument using an R9.4.1 flow cell (FLO-PRO002), with a run duration of 72 hours. Basecalling was performed using Guppy v7.1.4 with the ONT dna_r9.4.1_promethion_384 model, using MinKNOW v23.07.12.

### Data processing and bioinformatic analysis of Oxford Nanopore data

ONT base-called reads were processed using *epi2me-labs/wf-single-cell v0.2.8* analysis pipeline, with the following settings: “*--kit_name 3prime --kit_version v3 --expected_cells 4800*”. After stranding and adapter trimming, reads were mapped against Human Genome reference GRCh38 and GENCODE v32 (Ensembl 98) annotation using minimap2^37^. Isoforms were formed and refined using stringtie2 ^38^ with long read option ‘-L’. Cell barcodes were compared against 10X barcode whitelist, and corrected to remove non-cells. UMIs were clustered based on genomic location and corrected using Levenshtein distance. Corrected barcodes, UMIs, and stringtie2 isoforms were tagged in the BAM file.

The tagged BAM generated with wf-single-cell was used to run Isoquant (v3.4.1)^39^ using parameters recommended for ONT: --splice_correction_strategy conservative_ont --model_construction_strategy sensitive_ont. SQANTI3 ^40^ was run to classify isoforms into structural categories and filter out novel isoforms that are potentially intra-priming and template chatswitching artefacts. The default filter parameters were as follows: a novel isoform was kept if all of its splice junctions were either canonical or had more than 3 short reads spanning it. UMIs were deduplicated with UMI-Tools ^41^ and the resulting BAM was used to quantify the filtered transcripts per cell with IsoQuant without model construction using --read_group tag:CB --transcript_quantification unique_only --gene_quantification unique_only. The entries in the expression matrices generated by IsoQuant were originally indexed by transcriptID or geneID. To facilitate downstream analysis, we converted these to gene symbols, using a lookup table derived from the GRCh38-2024-A annotation (refdata-gex-GRCh38-2024-A). Novel genes not present in the reference were retained.

### Jumpcode CRISPRClean treatment

Aliquots of amplified cDNA from PBMC Samples 1 and 2 were (15 μL) was used as input for depletion with the CRISPRclean Single Cell RNA Boost Kit (Jumpcode Genomics, Cat. No: 10180). Size selection clean-ups were performed using SMRTbell Clean Up Beads (PacBio, Cat. No: 102-158-300). The starting material and post-depletion recovery were quantified using a Qubit 1.0 fluorometer with the High Sensitivity DNA Qubit assay (Invitrogen). Following depletion, the samples underwent MAS-seq library preparation.

### PacBio MAS-Seq library preparation and sequencing of PBMC cDNA

Aliquots of amplified cDNA from PBMC Samples 1 and 2 (15ng) were used to prepare MAS-seq (Multiplexed Arrays Sequencing) libraries using the PacBio MAS-Seq for 10X Single Cell 3’ Kit (PacBio, PN: 102-407-900). For each sample, two MAS-seq libraries were prepared: one from untreated cDNA and one from cDNA treated with Jumpcode CRISPRclean, resulting in a total of four MAS-seq libraries. Additional reagents used included MAS Capture Beads (Cat. No: 102-428-400) and SMRTbell Cleanup Beads (Cat. No: 102-158-300). Libraries were prepared following the manufacturer’s protocol (Procedure & Checklist 102-678-600, Revision 002).

This involved enrichment of biotin-tagged amplified cDNA using streptavidin-coated MAS Capture Beads to remove TSO artifacts, followed by 16 parallel PCR reactions per sample using premixed MAS primers to amplify cDNA with programmable sequences at both ends. The amplified cDNA was then assembled into linear arrays of transcripts (“segments”) with programmable ends before final SMRTbell library preparation.

A Qubit 1.0 fluorometer with the High Sensitivity DNA Qubit assay (Invitrogen) was used to quantify total yield at each stage of library preparation. The size distributions of the final SMRTbell libraries were assessed using the Femto Pulse System with the Genomic DNA 165 Kit (Agilent Technologies). All MAS-seq libraries were sequenced on both the PacBio Sequel IIe and PacBio Revio platforms, resulting in a total of eight sequencing runs.

### PacBio Sequel IIe

The loading calculations for the MAS-Seq libraries were prepared using the PacBio SMRTlink Binding Calculator v11.1.0 and prepared for sequencing applicable to the library type. Sequencing primer 3.2 was annealed to the libraries and complexed to the sequencing polymerase with the Sequel II binding kit 3.2 (PacBio, 102-333-300). Calculations for primer to template and polymerase to template binding ratios were kept at default values for the library type. Sequencing internal control complex 3.2 (PacBio, 102-249-600) was spiked into the final complex preparation at a standard concentration before sequencing for all preparations. The sequencing chemistry used was Sequel® II Sequencing Plate 2.0 (PacBio®, 101-820-200) and the Instrument Control Software v11.0.1. Each MAS-Seq library was sequenced on the Sequel IIe instrument with one Sequel II SMRT®cell 8M cell. The parameters for sequencing were adaptive loading, 30-hour movie, 2-hour pre-extension time, 100pM on plate loading concentration.

### PacBio Revio

The loading calculations for the MAS-Seq libraries were prepared using the PacBio SMRTlink Sample Setup Calculator v12.0 and prepared for sequencing applicable to the library type. Standard Revio Sequencing primer was annealed to the libraries and complexed to the sequencing polymerase with the Revio polymerase kit (PacBio, 102-739-100) at the standard ratio (1:1). Revio internal control complex (PacBio, 102-798-000) was spiked into the final complex preparation at a standard concentration before sequencing for all preparations. The sequencing chemistry used was Revio Sequencing Plate (PacBio®, 102-587-400) and the Instrument Control Software v12.0. Each MAS-Seq library was sequenced on the Revio instrument with one Revio 25M SMRTcell (PacBio®, 102-202-200). The parameters for sequencing were 100 minutes diffusion loading, 24-hour movie, 2-hour pre-extension time, 225-250pM on plate loading concentration.

### Data processing and bioinformatic analysis of PacBio data

The PacBio MAS-Seq HiFi reads were processed using SMRT-Link v12.0 pb_segment_reads_and_sc_isoseq analysis pipeline. Key input parameters included the MAS-Seq Adapter v1 (MAS16) for segmentation adapters, 10X Chromium Single Cell 3’ cDNA Primers, the 3M-february-2018-REVERSE-COMPLEMENTED cell barcode whitelist, and the Human Genome hg38 reference with Gencode v39 annotations for minimap2^37^ alignment.

HiFi reads were deconcatenated into segments, before trimming off poly-A tails and cDNA primer sequences. Cell barcodes were extracted and corrected against the 10X whitelist, with real cells identified using the barcode “knee” finding method. Unique molecular identifiers (UMIs) were clustered by cell barcode and deduplicated using the *iso-seq groupdedup* tool to correct for PCR artifacts. Reads were aligned and collapsed to form isoforms, then classified by *pigeon classify* into categories defined in SQANTI3 ^40^. Filtering was applied via *pigeon filter* to exclude isoforms derived from artifacts, intrapriming events, or those classified as low confidence. Final isoform- and gene-level expression matrices were generated using the *pigeon make-seurat* command using default parameters, including the “*--keep-ribo-mito-genes*” option, enabling the retention of mitochondrial and ribosomal gene counts.

### Long-read single cell gene- and isoform-expression data analysis

Single-cell RNA-seq data were processed and analysed using Seurat (v5.1.0) ^34^. For each sample, a Seurat object was created, retaining cells with a minimum of five detected features. Isoform-level expression data were incorporated as an additional assay within each object. Quality control filters were applied to exclude low-quality cells, removing those with RNA feature counts outside the 1st and 99th percentiles or mitochondrial gene content exceeding 15%.

Standard preprocessing steps included log-normalisation, identification of highly variable features, and dimensionality reduction using Principal Component Analysis (PCA). To correct for batch effects across samples, where applicable, integration was performed using canonical correlation analysis (CCA), and the integrated dataset was used for downstream analyses. UMAP was applied to the first 20 principal components for visualisation, and clustering was performed using the Louvain algorithm at a resolution of 0.8, with gene-level cluster annotations stored as metadata.

For isoform-level analyses, the isoform assay was set as the default and log-normalised. Seurat objects were split by sample, and variable features were identified within each sample, with the intersection of variable features across samples used for downstream analysis. Data scaling and PCA were performed on the isoform assay, followed, where applicable, by batch correction using Harmony, grouping by sample of origin. UMAP was applied to the Harmony-corrected space for dimensionality reduction, and clustering was performed using a shared nearest neighbour (SNN) graph at a resolution of 0.8, with isoform-level or combined (gene- and isoform-level) cluster annotations stored as metadata.

A clustering resolution of 0.8 was used consistently across gene- and isoform-level analyses to facilitate direct comparisons. This resolution was selected after testing multiple values (0.2, 0.5, 0.8, 1.0, 1.2) and was found to provide the most accurate identification of expected PBMC cell populations.

Cell type annotations were assigned based on canonical marker gene expression cross-validated using the automated annotation tools ScType ^35^ and SingleR ^36^, followed by manual refinement.

### Immune receptor reconstruction (TRUST4)

Immune receptor repertoires in αβ/γδ T cells and B cells were reconstructed for each PBMC sample across all sequencing approaches (except for ONT, where only Sample 1 was available) using the TRUST4 pipeline^15^. In each case, the aligned bam files were used as input. The appropriate human hg38 reference file (--ref) IGMT+C.fa and database file (-f) bcrtcr.fa were generated with perl scripts provided by the authors (https://github.com/liulab-dfci/TRUST4, accessed 30/03/2023), using hg38 data downloaded from IGMT (https://www.imgt.org/ accessed 30/03/2023) following Song *et al* ^15^. In each case, the single-cell barcode field in each bam file was specified by --barcode CB and the UMI field with --UMI XM for PacBio and --UMI UB for ONT. The resulting data (barcode_report.tsv) were parsed to extract proportions of B, αβ-T and γδ-T cells with successfully reconstructed primary chain-1 and chain-2 for each sequencing approach and plotted using custom python scripts (see code availability). Clonal representation across long-read approaches was visualised using the R package scRepertoire v2.0.0 ^42^.

## DATA ACCESSIBILITY STATEMENT

Analysis scripts are available from https://github.com/TGAC/LR_scRNAseq_Scoones_etal. Data will be made available upon publication in appropriate repositories (ENA / Zenodo).

## Acknowledgements

The authors acknowledge support from an AIRC/CRUK/FC AECC Accelerator Award ’Single Cell Cancer Evolution in the Clinic’, A26815 (AIRC number program 2279) and Biotechnology and Biological Sciences Research Council (BBSRC), part of UK Research and Innovation (UKRI), Core Capability BB/CCG1720/1 and BB/CCG2220/1 and National Capability (BBS/E/T/000PR9816), Earlham Institute Strategic Programme Grant Cellular Genomics BBX011070/1 grants at the Earlham Institute (EI). WH was also supported by UKRI grant EP/X035913/1, and Innovate UK grant 10098097 and DW by BBSRC grant BB/V016156/1. SK, CU and LP were supported by the BBSRC funded Norwich Research Park Biosciences Doctoral Training Partnership grant BB/T008717/1 and AS by BB/M011216/1. Single-cell and sequencing work was delivered through Transformative Genomics, the BBSRC funded National Bioscience Research Infrastructure (BBS/E/ER/23NB0006) at EI by members of the Technical Genomics and Core Bioinformatics Groups. We also thank the Laboratory Managers and Research Computing Groups at EI who manage and deliver High Performance Computing at EI alongside the physical HPC infrastructure and data centre delivered via the Norwich Bioscience Institutes (NBI) Computing infrastructure for Science (CiS) group. A.P.C. is a recipient of a Medical Research Council (MRC) Career Development fellowship (grant no. MR/V010182/1).

## Supplementary Material

**Figure S1.**
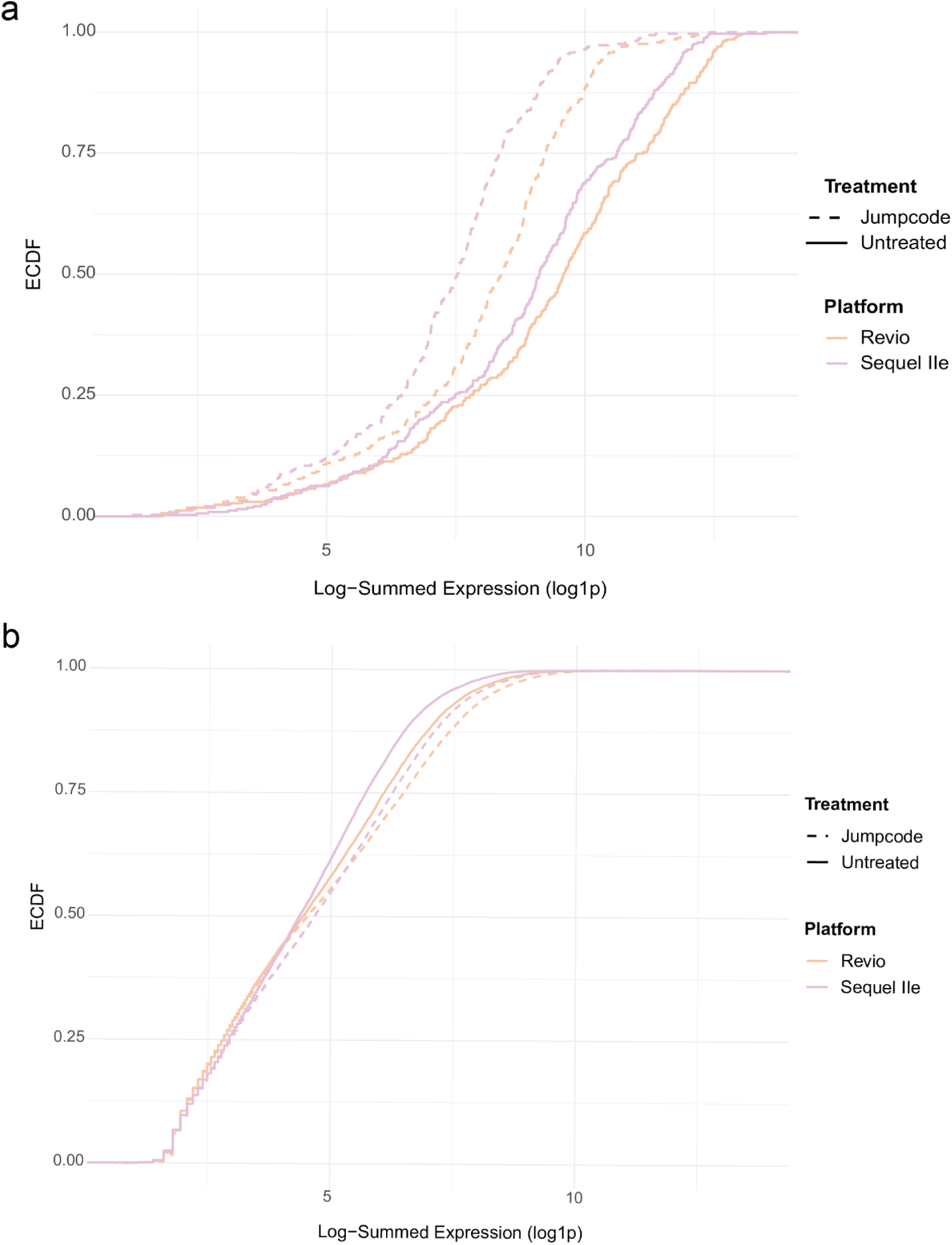
ECDFs of Summed Gene Expression After Jumpcode Treatment Empirical cumulative distribution functions (ECDFs) showing log₁-transformed summed expression of panel and non-panel genes across Sequel and Revio platforms, with and without Jumpcode treatment.For panel genes, Jumpcode treatment shifted the distributions towards lower summed expression values, consistent with targeted transcript depletion. In contrast, non-panel genes exhibited minimal shifts in distribution following treatment, supporting the specificity of the Jumpcode depletion approach.

**Supplementary Figure S2.**
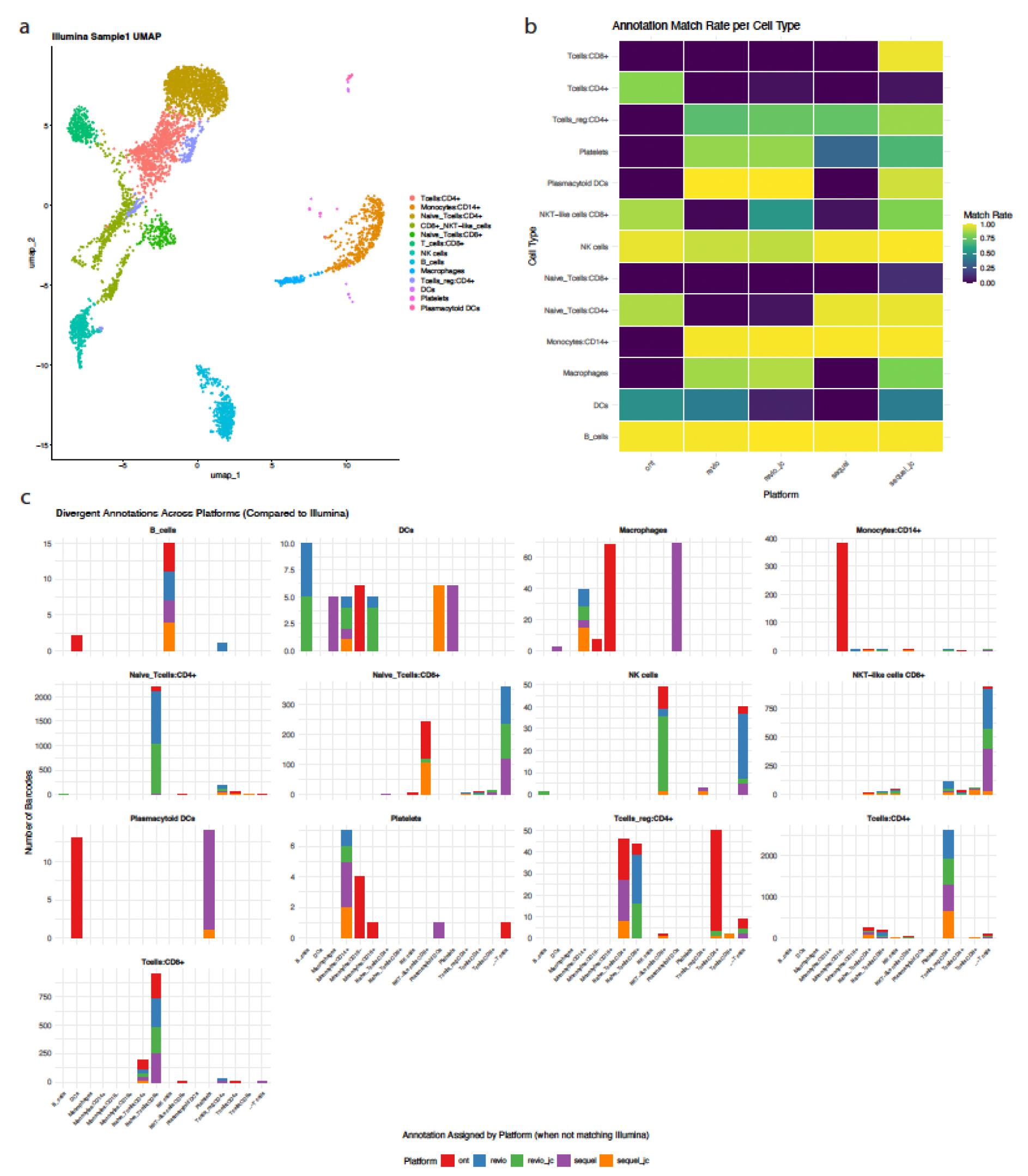
Platform comparison of single-cell annotations using Illumina labels as a reference. (a) UMAP projection of PBMCs annotated using short-read Illumina data, coloured by cell type (b) Heatmap showing the number of barcodes per platform with annotations matching Illumina, across cell types. (c) Stacked bar plots showing the distribution of alternative annotations for each Illumina-annotated cell type. Each panel represents one true (Illumina-assigned) label, and bars indicate how cells were reclassified across platforms when their annotation did not match Illumina. X-axis: number of barcodes; Y-axis: alternative label assigned.

**Supplementary Figure S3:**
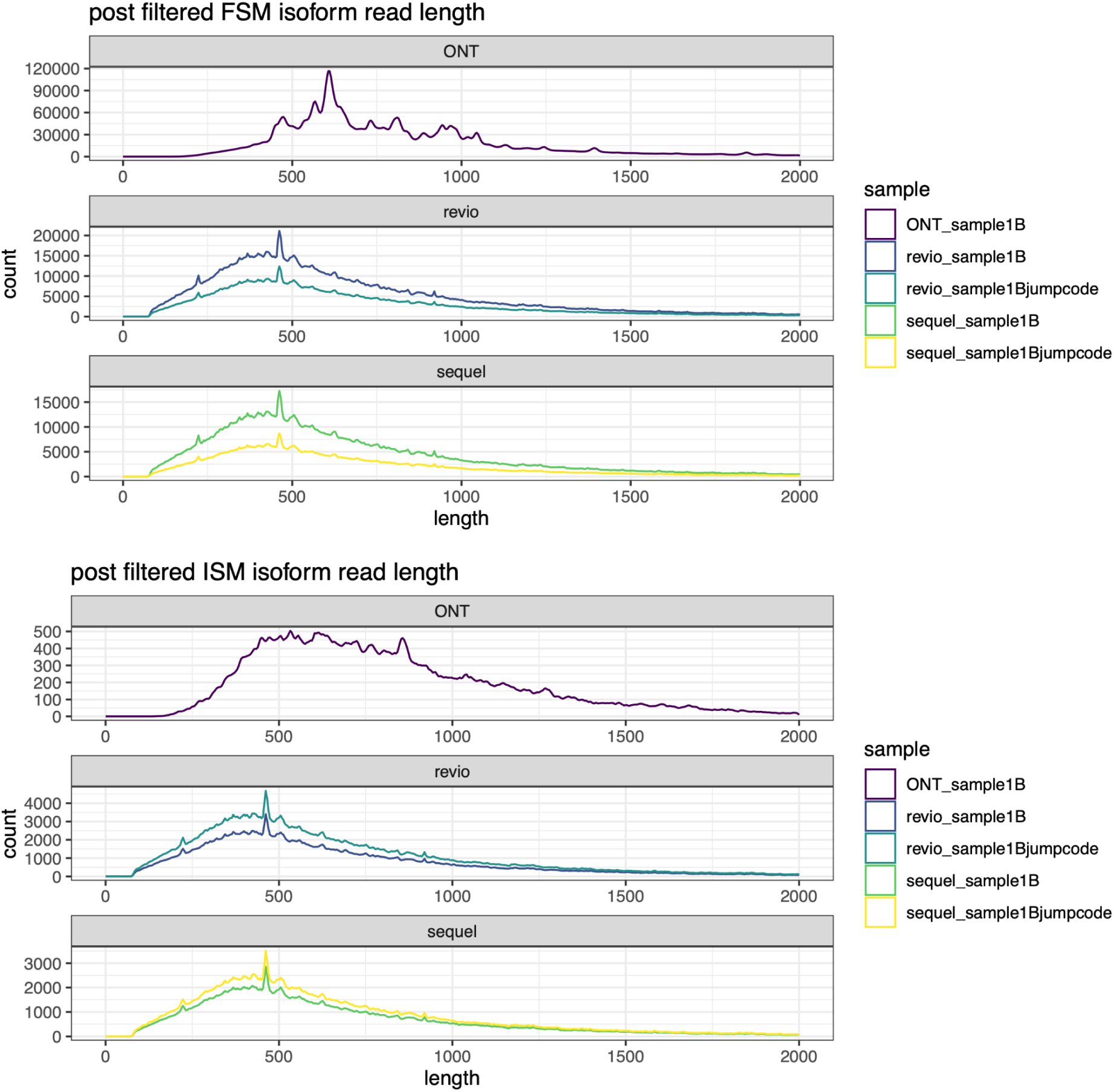
Length distribution for deduplicated reads associated with PacBio and ONT FSM / ISM isoforms which were isoform filtered. Reads are shorter than normal iso-seq reads.(Note the different range on y-axis)

**Supplementary Figure S4:**
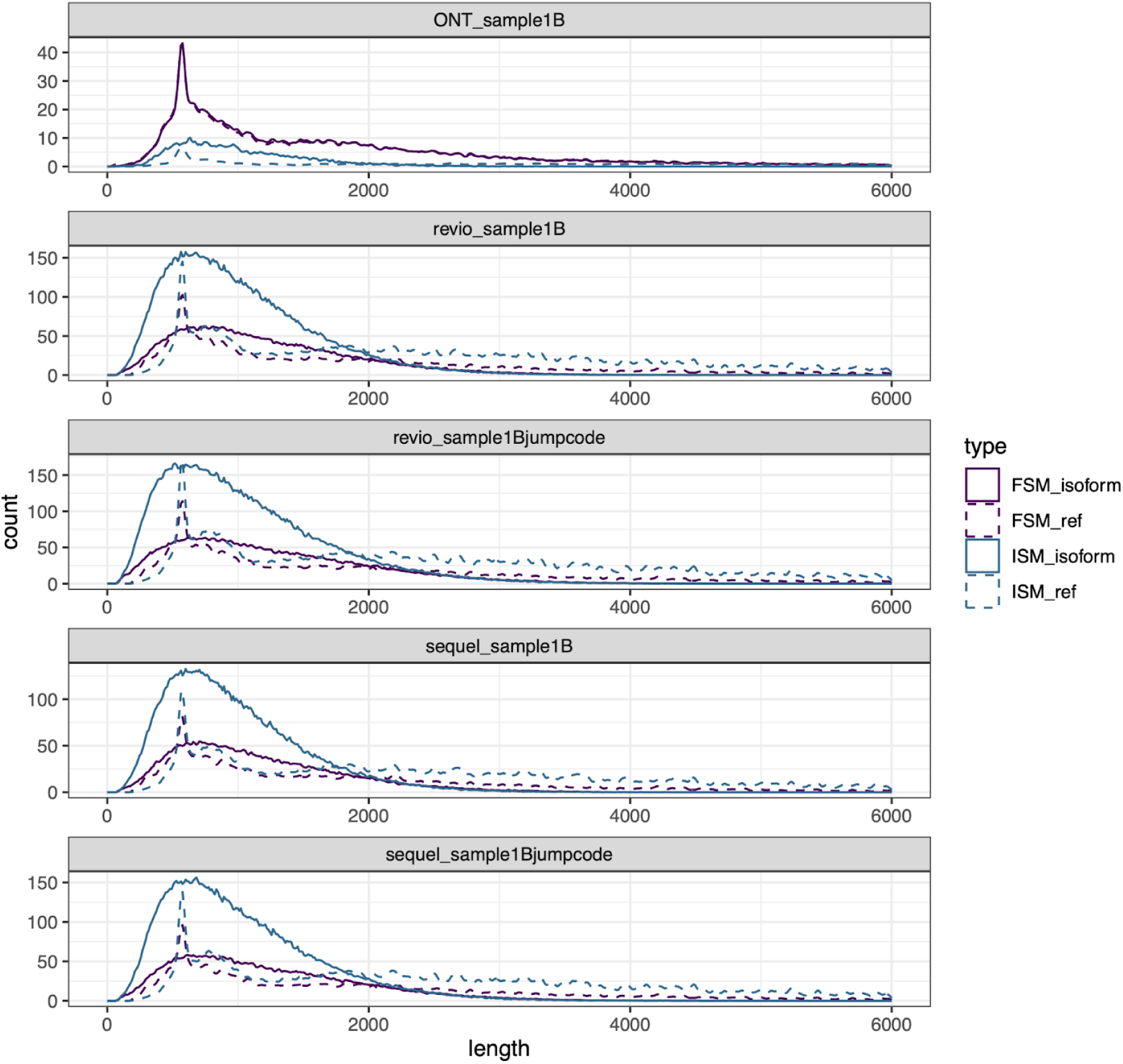
Post-filtered FSM / ISM isoform length versus transcript reference. Comparing ONT & PacBio isoform length post pigeon-filtering to its reference.

**Supplementary Figure S5:**
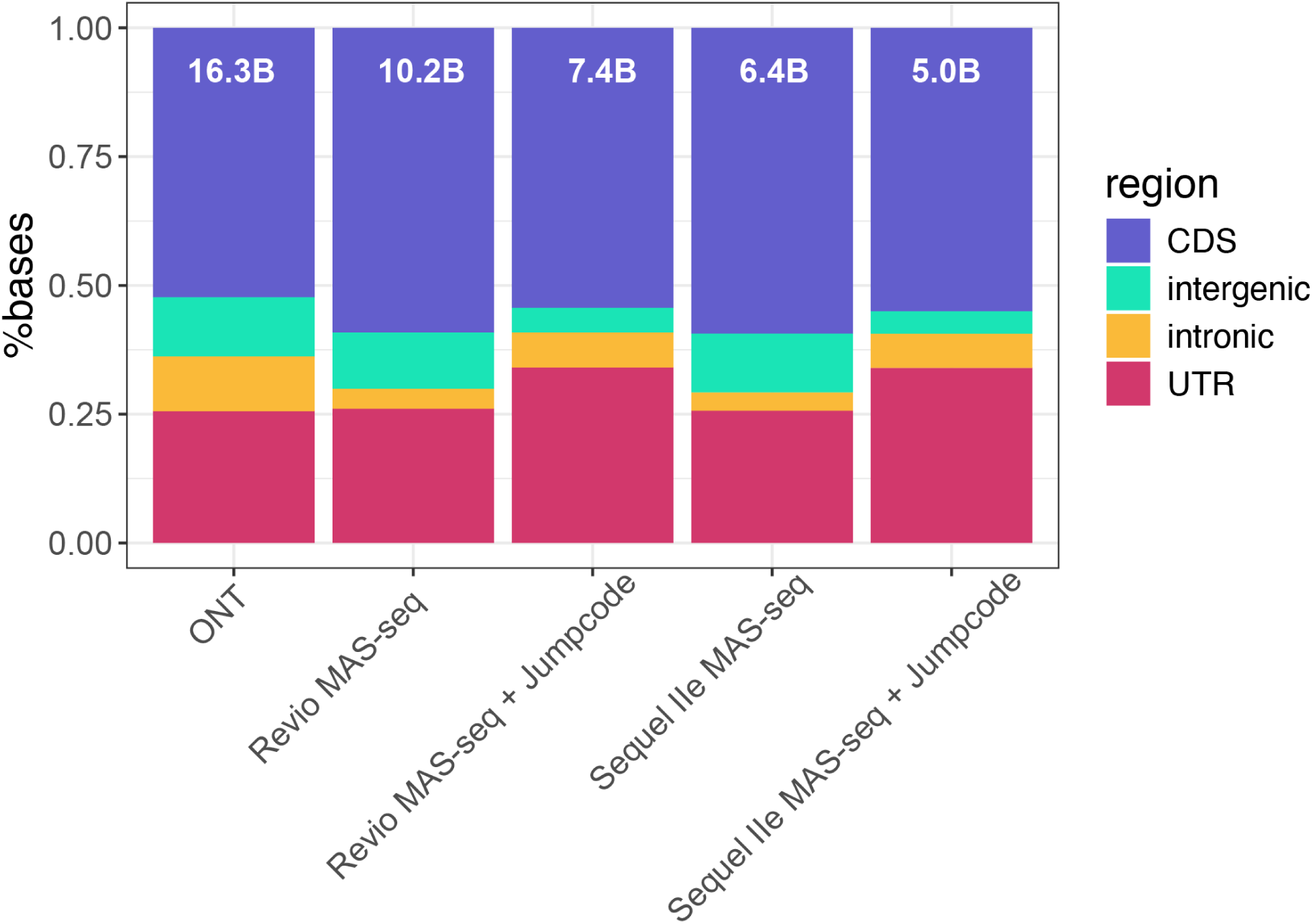
Distribution of bases for post-isoform filtered deduplicated reads that align to reference genome. Column number is the total bases aligned per region.

**Supplementary Figure S6:**
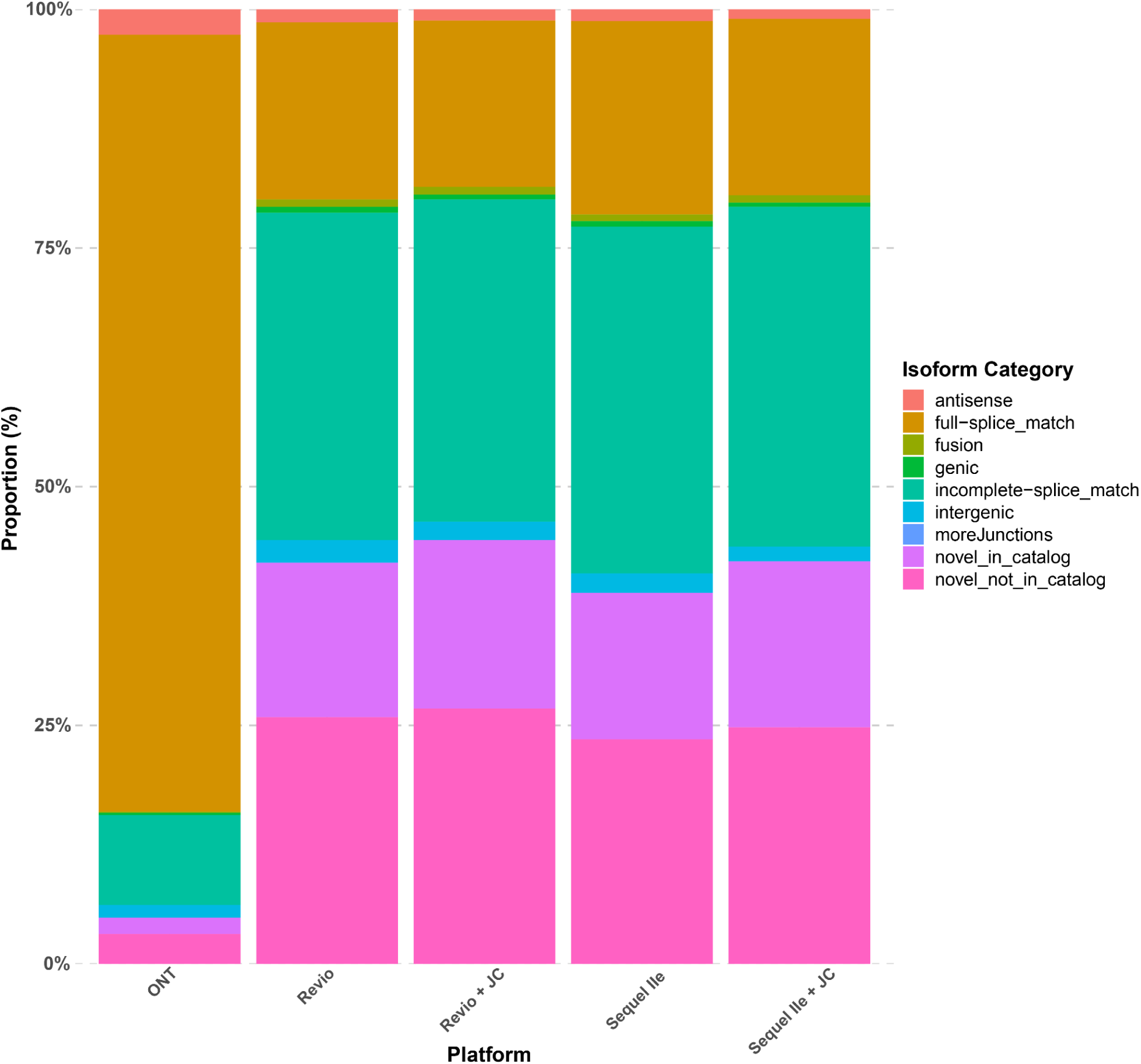
Proportion of isoforms per SQANTI structural category across long-read methods & platforms.

**Supplementary Figure S7:**
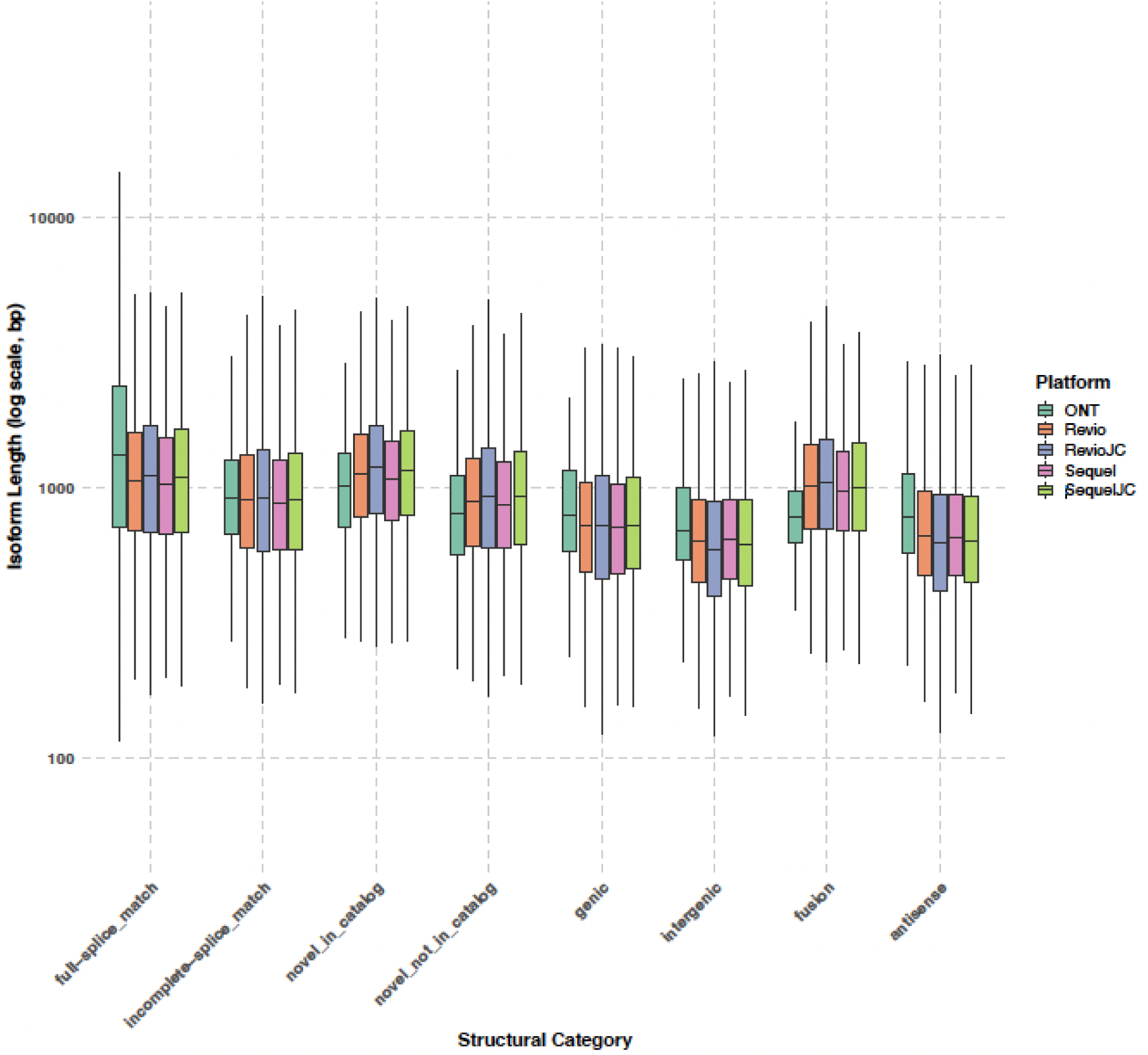
Distribution of isoform length for each SQANTI structural category across long read methods.

**Supplementary Figure S8.**
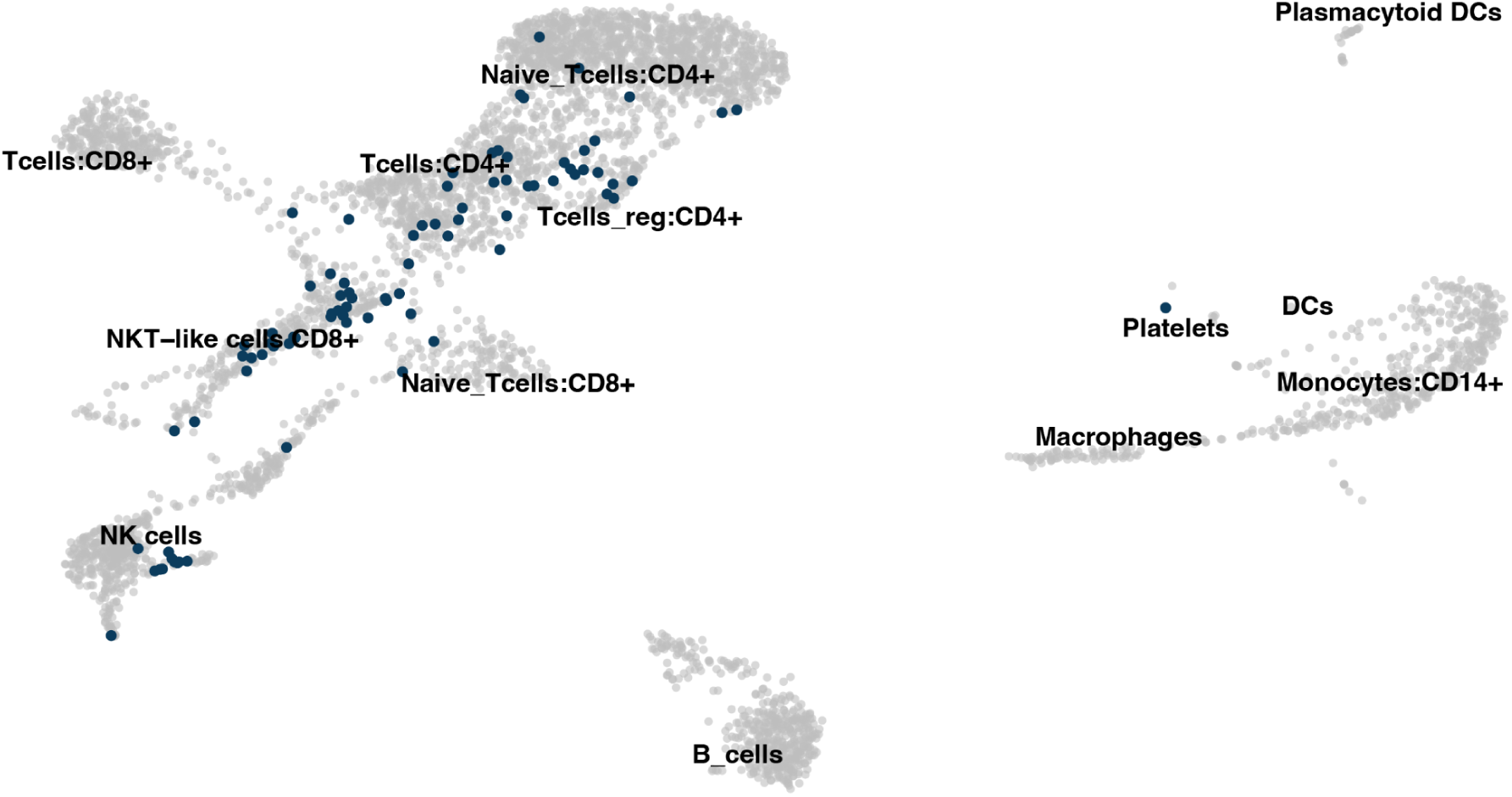
UMAP of Illumina short-read data with only gamma delta T cells as assigned by TRUST4 coloured (blue).

**Supplementary Figure S9.**
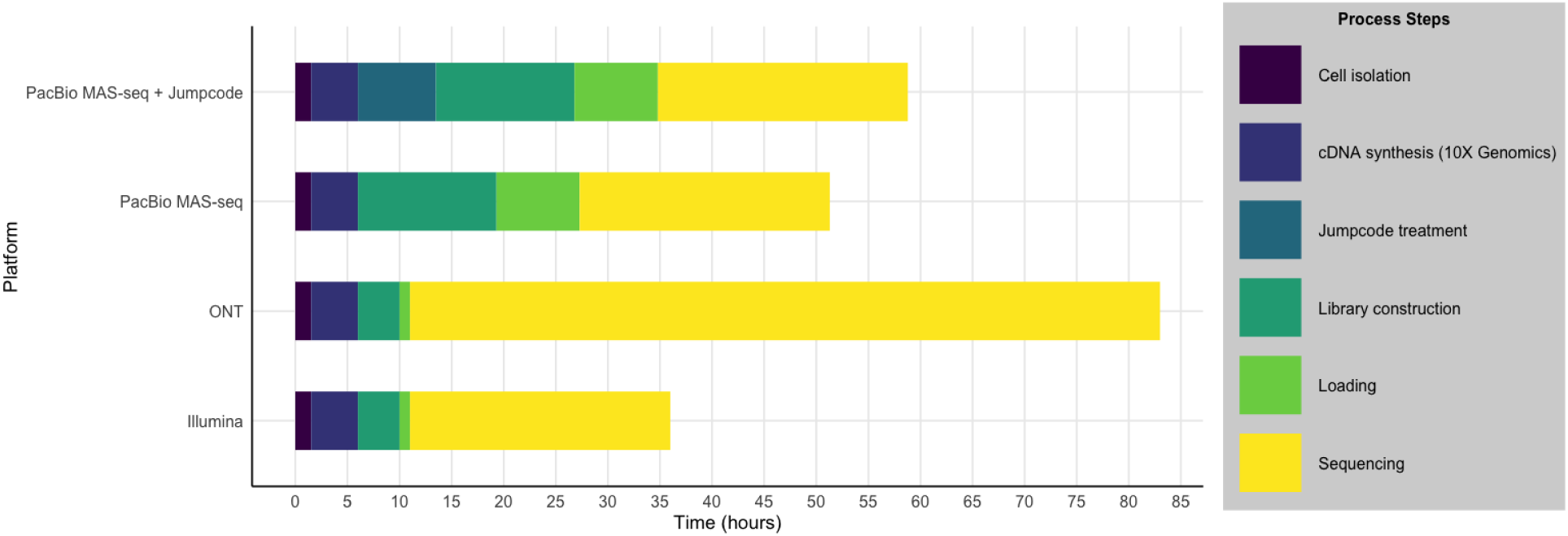
Stacked bar plot showing current processing time from cell isolation to completion of sequencing for each of the platforms implemented in this study: Illumina, ONT (Oxford Nanopore Technologies), PacBio MAS-seq, and PacBio MAS-seq with Jumpcode depletion. Segment colours represent specific steps in the experimental workflow, including cell isolation, cDNA synthesis (10X Genomics), Jumpcode treatment (if applicable), library construction, loading, and sequencing.

